# *In vitro* characterisation and neurosteroid treatment of an N-Methyl-D-Aspartate receptor antibody-mediated seizure model

**DOI:** 10.1101/2020.12.22.423962

**Authors:** Sukhvir K Wright, Richard E Rosch, Max A Wilson, Manoj A Upadhya, Divya R Dhangar, Charlie Clarke-Bland, Tamara T Wahid, Sumanta Barman, Norbert Goebels, Jakob Kreye, Harald Prüss, Leslie Jacobson, Danielle S Bassett, Angela Vincent, Stuart D Greenhill, Gavin L Woodhall

## Abstract

Seizures are a prominent feature in N-Methyl-D-Aspartate receptor antibody (NMDAR-Ab) encephalitis, a distinct neuro-immunological disorder in which specific human autoantibodies bind and crosslink the surface of NMDAR proteins thereby causing internalization and a state of NMDAR hypofunction. To further understand ictogenesis in this disorder, and to test a novel treatment compound, we developed an NMDAR-Ab mediated rat seizure model that displays spontaneous epileptiform activity *in vivo* and *in vitro*. Using a combination of electrophysiological and dynamic causal modelling techniques we show that, contrary to expectation, reduction of synaptic excitatory, but not inhibitory, neurotransmission underlies the ictal events through alterations in the dynamical behaviour of microcircuits in brain tissue. Moreover, *in vitro* application of an NMDAR-specific neurosteroid, pregnenolone sulfate, that upregulates NMDARs, reduced established ictal activity. This proof-of-concept study highlights the complexity of circuit disturbances that may lead to seizures and the potential use of receptor-specific treatments in antibody-mediated seizures and epilepsy.

## Introduction

Both *in vivo* and *in vitro* studies have clearly established that the pathogenic action of antibodies in N-Methyl-D-Aspartate receptor antibody (NMDAR-Ab) encephalitis is to bind to the surface of the NMDARs, cross-link the proteins and cause NMDAR internalization, resulting in a state of NMDAR hypofunction (1–3). As a result, the mainstay of treatment is immunotherapy that aims to reduce the levels of circulating neuronal autoantibodies or halt their production (4, 5). These treatments can be slow to work, and carry risk of significant sideeffects as the healthy immune system is also compromised (6). It is clear that the reduction of NMDARs in NMDAR-Ab encephalitis causes a myriad of severe neurological symptoms, and further that reversal of this pathological effect in the acute and even chronic stages of the disease has the potential to improve clinical outcomes (7–9). However, as yet, there are no immune-sparing treatments that ameliorate the synaptic and circuit effects (psychosis, ictogenesis) caused by the NMDAR hypofunction (10, 11). The therapeutic potential of a receptor-specific treatment would be particularly beneficial in children who are affected during crucial developmental stages, marked by neuronal network plasticity including that of the NMDAR subunits (12, 13). Indeed, studies have found that after recovery children have persistent cognitive problems and fatigue, resulting in lower academic achievement and poorer quality of life (14, 15).

While neuropsychiatric features are the most common presenting symptom of NMDAR-Ab encephalitis, acute symptomatic seizures and encephalopathic electroencephalogram (EEG) changes are essential clinical and investigative features in patients of all ages (16). Seizures present as part of the disease course in >80% of patients and EEG changes are seen in >95%, more commonly than Magnetic Resonance Imaging (MRI) brain changes, as noted repeatedly in cohort and case studies (4, 5, 17). Despite the symptom predominance of seizures in humans, producing reliable and consistent spontaneous seizure animal models of this disorder using passive transfer of human-derived antibodies is challenging (18–20). In our previous passive transfer mouse model, we demonstrated increased *in vivo* seizure susceptibility (and therefore network hyperexcitability) 48 hours after intracerebroventricular NMDAR-Ab injection compared to controls, however spontaneous seizures were not seen (18, 21).

Importantly, seizures within the context of NMDAR-Ab encephalitis do not respond well to standard anti-seizure medications, as seen in other forms of antibody mediated encephalitis (22–24). Given the specific involvement of the excitatory NMDAR-receptor, it is tempting to speculate that the pathophysiology of these epilepsies diverges from a simple ‘excitation-inhibition’ imbalance and that an NMDAR-specific treatment may be more effective in minimizing acute symptoms such as seizures, and could potentially be given alongside immunotherapy to achieve more rapid symptomatic relief.

This study aimed to understand the causative changes in synaptic physiology contributing to NMDAR-Ab mediated seizures, and to explore the unmet clinical need for receptor-specific treatment. We developed a novel juvenile Wistar rat model of NMDAR-Ab encephalitis, in which NMDAR-Abs derived from human patients were used to induce a robust and repeatable seizure phenotype. Combining *in vitro, in vivo*, and *in silico* approaches we have demonstrated that, counterintuitively, the epileptic dynamics emerge from reductions in excitatory neurotransmission caused by NMDAR-Abs, and provide proof-of-concept *in vitro* evidence of a NMDAR-Ab specific neurosteroid rescue treatment both in the rodent model, and in human epileptic brain tissue.

## Results

### NMDAR-Abs bind specifically to Wistar rat brain tissue

All three NMDAR-Ab preparations derived from patients with the disease bound specifically to rodent hippocampus (Supplementary Figure 1) confirming previous studies in mice (3, 8, 18). The florescence intensities of NMDAR-Ab injected and infused brain slices were significantly higher than those measured after injection or infusion of control human IgG or non-brain reactive monoclonal antibodies (mGo53, 12D7) (Supplementary Figure 1a-g).

### NMDAR-Abs cause spontaneous epileptiform activity *in vitro*

In order to characterize NMDAR-Ab induced abnormalities of synaptic transmission and circuit dysfunction, we first examined the effects of the antibody preparations on brain-slices *in vitro*.

At 48 hours after intracerebroventricular (ICV) injection, juvenile rats injected with NMDAR-Abs exhibited sustained and repetitive hyperexcitable behavior including myoclonic twitches, jerks and jumps (Supplementary Video 1). Animals were sacrificed at this time-point and brain slices were prepared for *in vitro* electrophysiology studies. These hippocampal local field potential (LFP) recordings showed significantly more frequent spontaneous epileptiform activity in NMDAR-Ab injected slices compared to slices prepared from control-Ab injected animals (Figure 1a-d). In the NMDAR-Ab injected animals the number of interictal events was higher in the CA3 hippocampal region compared to CA1, with reduced interevent interval (IEI) between spikes (Figure 1e, f).

**Figure 1.**
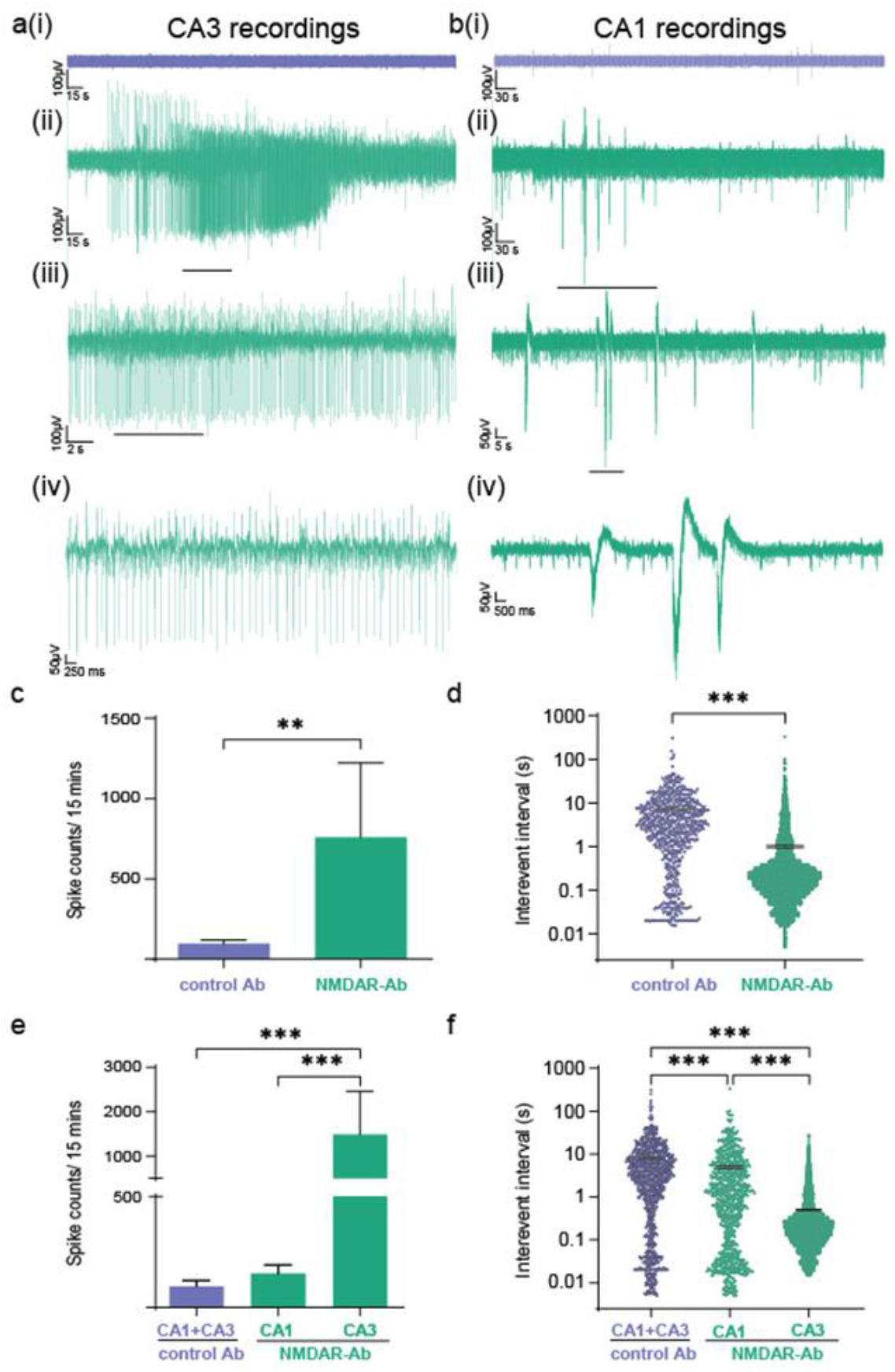
Hippocampal local field potential recordings *in vitro* 48 hours after intracerebroventricular injection of NMDAR-Abs show spontaneous epileptic activity, highest in the CA3 region. **a** Example traces of local field potential brain slice recordings from control (ai) and NMDAR-Ab (aii-iv) injected animals after 48 hours from the CA3 region. Scale bar 100μV vs. 15 sec (i and ii), 100μV vs. 2 sec (iii) and 50μV vs. 250 msec (iv). **b** Example traces of local field potential brain slice recordings from control (bi) and NMDAR-Ab (bii-iv) injected animals after 48 hours from the CA1 region. Scale bar 100μV vs. 30 sec (I and ii), 50μV vs. 5 sec (iii) and 50μV vs. 500 msec (iv). **c** The number of spikes (interictal events) per 15 minutes in the NMDAR-Ab slices compared to controls (Control (n=8) vs. NMDA (n=11); p<0.01, Mann-Whitney). **d** The intervent interval (IEI) in control Ab slices compared to NMDAR-Ab injected slices (Control (n=8) vs. NMDA (n=11); p<0.001, Mann-Whitney). **e** The number of spikes (interictal events) in the CA3 region of the NMDAR-Ab hippocampal slices as compared to CA1 (NMDA CA1 (n=6) vs. NMDA CA3 (n=5); p<0.001, Mann-Whitney). **f** Comparison of IEI between the CA3 and CA1 region in NMDAR-Ab hippocampal slices (NMDA CA1 (n=6) vs. NMDA CA3 (n=5); p<0.001, Mann-Whitney). Measurements expressed as mean ± standard error of the mean (SEM).

To determine the synaptic changes contributing to this apparent circuit hyperexcitability, whole-cell patch clamp recordings were made from pyramidal cells in the CA3 region of the hippocampus (Figure 2a; cell morphology confirmed on *post-hoc* analysis, Supplementary Figure 1h). There was a reduction in the frequency and amplitude of spontaneous excitatory post-synaptic current (sEPSC) recordings (Figure 2b-d) and a significant shift towards a faster decay time indicating loss of NMDAR-mediated events (Figure 2e, f). By contrast, no changes were seen in the frequency or amplitude of spontaneous inhibitory post-synaptic currents (sIPSCS) or miniature IPSCs (Figure 2g-l). Together these results indicate that local hippocampal networks are hyperexcitable when recorded *in vitro* 48 hours after a single ICV injection of NMDAR-Abs *in vivo*. This circuit hyperexcitability is paradoxically associated with a reduction of excitatory neurotransmission in CA3 induced by the NMDAR-Abs.

**Figure 2.**
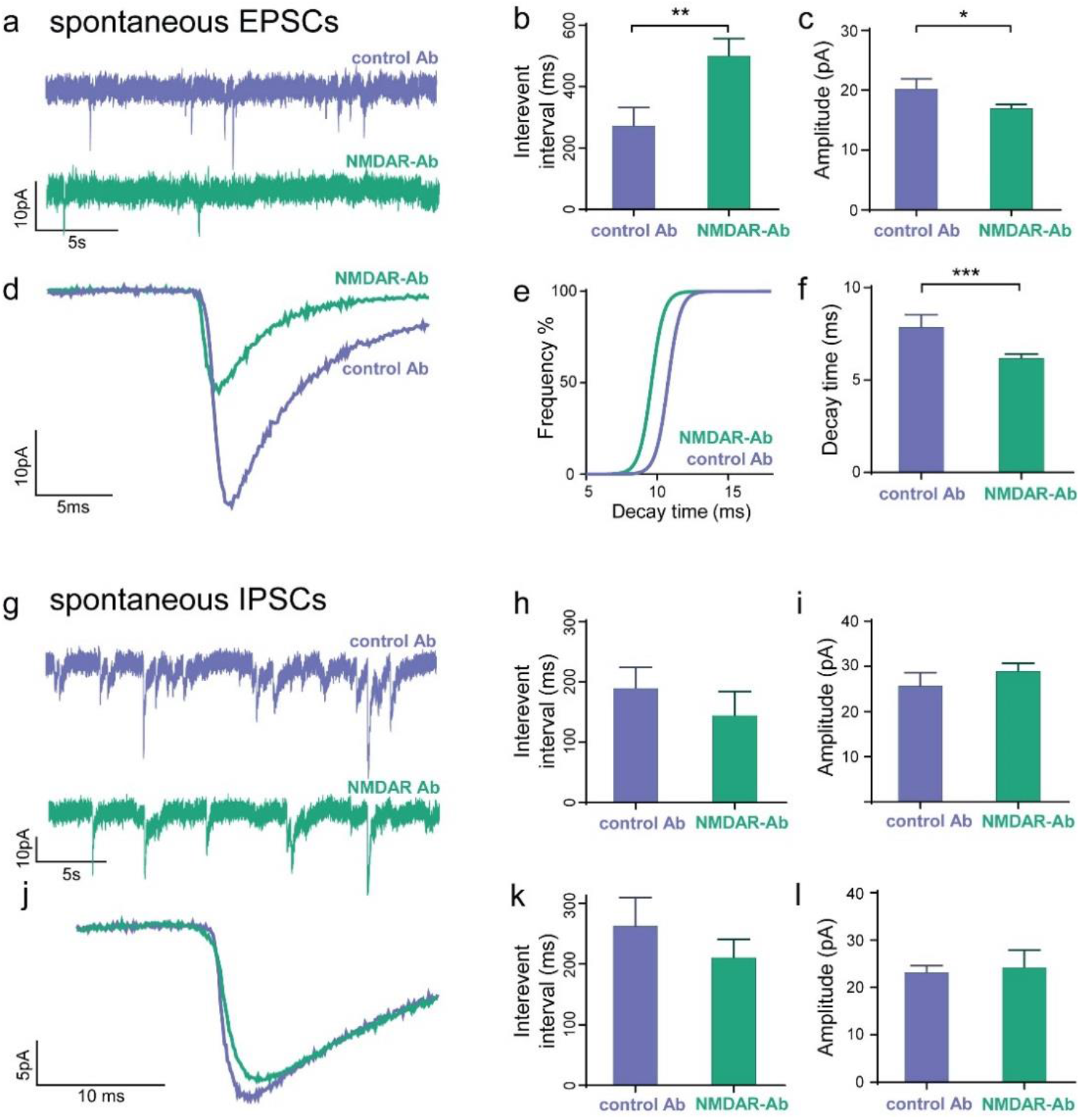
Whole-cell patch-clamp recordings from hippocampal CA3 pyramidal cells *in vitro* 48 hours after intracerebroventricular injection of NMDAR-Abs show a reduction in excitatory but not inhibitory synaptic neurotransmission. **a** Representative spontaneous EPSC (sEPSC) whole-cell patch clamp recordings 48 hours after injection with NMDAR-Abs (lower trace) or control Abs (upper trace). Scale bar 10pA vs. 5s. **b** The interevent interval (IEI) of sEPSCs recorded from CA3 pyramidal cells in NMDAR-Ab injected rodent slices (n=18 cells) compared to control Ab (n=8 cells) injected slices (p=0.003, Mann-Whitney). **c** The amplitude of the sEPSCs in CA3 pyramidal cells recorded from NMDAR-Ab injected slices (n=18) compared to controls (n=8) (p=0.04; Mann-Whitney). **d** Representative average sEPSCs recorded from NMDAR-Ab injected slices as compared to control-Ab injected slices. Scale bar 10pA vs. 10ms. **e** Cumulative frequency plot showing reduced decay time of sEPSCs recorded from NMDAR-Ab slices as compared to the controls (n=7 cells in each group). **f** Decay time of the sEPSCs in the NMDAR-Ab injected rodent slices as compared to the controls (n= 7 cells in each group; p= <0.001, Mann-Whitney). **g** Representative whole-cell patch clamp recording of spontaneous IPSC (sIPSC) from CA3 pyramidal cells after injection of NMDAR- or control-Abs. Scale bar 10pA vs. 5s. **h** The intervent interval of the sIPSCs recorded from CA3 pyramidal cells in control and NMDAR-Ab injected slices (n=6 cells in each group; p=ns, Mann-Whitney). **i** The amplitude of sIPSCS recorded from the control and NMDAR-Ab injected slices (n=6 cells in each group; p=ns, Mann-Whitney). **j** Representative average sIPSCs recorded from NMDAR-Ab injected slices as compared to control-Ab injected slices. Scale bar 5pA vs. 10ms. **k** The interevent interval of miniature IPSCs recorded (in the presence of TTX 1μM) from pyramidal cells in CA3 in control and NMDAR-Ab injected slices (n=6 cells in each group; p=ns, Mann-Whitney). **l** The amplitude of miniature IPSCs recorded (in the presence of TTX 1μM) from pyramidal cells in CA3 in control and NMDAR-Ab injected slices (n=6 cells in each group; p=ns, Mann-Whitney). Measurements expressed as mean (M) ± standard error of the mean (SEM).

### NMDAR-Abs cause spontaneous epileptic seizures *in vivo*

To demonstrate ictogenesis *in vivo*, 7-day osmotic pumps were used to deliver NMDAR-Abs or control-Abs into the lateral cerebral ventricles of juvenile Wistar rats (n=6 for each group). Following the observations above, a depth electrode was placed in the CA3 region. Spontaneous epileptiform events were observed in the NMDAR-Ab infused animals *in vivo* and evident in the EEG tracings (Figure 3a, Supplementary Video 2a-c). A significant increase in EEG coastline of the NMDAR-Ab animals compared to control-Ab treated reflected this increase in occurrence of epileptiform high amplitude events (Figure 3b). Using automated seizure detection (18, 25), we observed a significant increase in the number of ictal events per hour in the NMDAR-Ab animals compared to controls. The maximum ictal event frequency was seen in NMDAR-Ab animals during Day 2 of the infusion, but it remained significantly elevated compared to controls until the end of the recording and infusion (Figure 3c-e). The EEG of NMDAR-Ab animals exhibited increased power in all frequency bands measured compared to control-Ab animals (1-4Hz, 4-8Hz, 8-12Hz, 12-30Hz, 30-50Hz, 50-70Hz, 70-120Hz, and 120-180Hz; p= <0.0001, Mann-Whitney). The greatest increase in power, seen in the 12 to 30 Hz range (676 ±151 vs 3195 ± 102, n=6; p= <0.0001, Mann-Whitney), was similar to the typical excessive beta activity observed in EEGs of NMDAR-Ab patients (14 to 20 Hz) (26).

**Figure 3.**
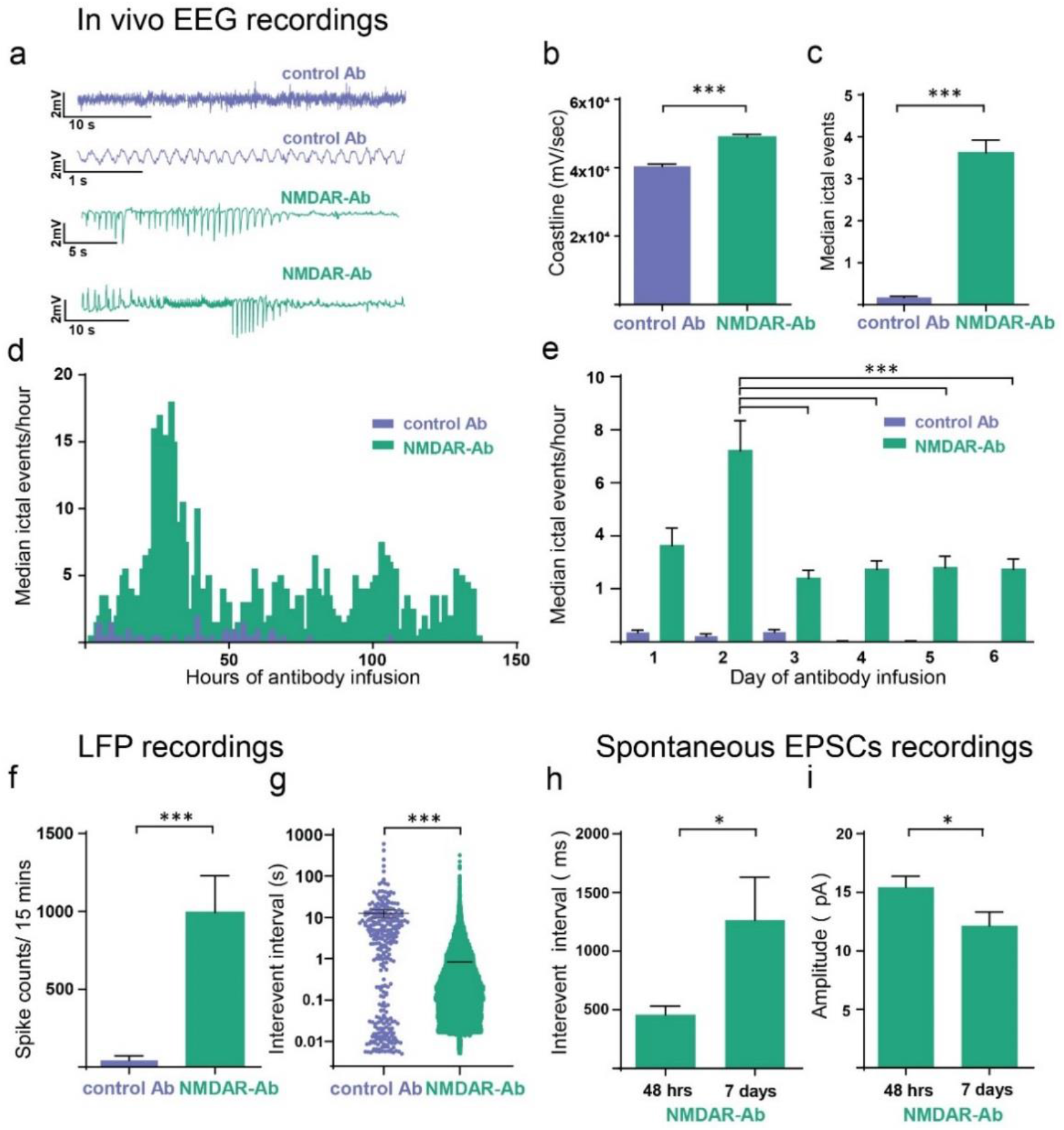
*In vivo* spontaneous epileptic EEG activity recorded in rodents infused with NMDAR-Abs. **a** Example EEG recordings from experimental rodents using wireless EEG transmitters with osmotic pumps *in situ* delivering control or NMDAR-Abs. **b** EEG coastline length (calculated per second for the total 7 day recording period for each animal) in NMDAR-Ab animals as compared to control animals (n=6 animals in each group; p<0.0001, Mann-Whitney). **c** The number of spontaneous ictal events/hour (mean median ± SEM) in the NMDAR-Ab animals compared to controls (p<0.001; Mann-Whitney).**d** Graph illustrating median number of ictal events over time of control and NMDAR antibody infusion. **e** Daily comparison of ictal events in the NMDAR-Ab and control-Ab animals (p=<0.001, two-way Anova with Bonferroni correction). **f** Ictal spike counts from hippocampal LFP measurements in brain slices prepared from NMDAR-Ab and control-Ab infused animals at the end of the *in vivo* EEG recording period (controls (n=7) vs. NMDA (n=38); p<0.05, Mann-Whitney. **g** Interevent interval from hippocampal LFP measurements in brain slices prepared from NMDAR-Ab and control-Ab infused animals at the end of the *in vivo* EEG recording period (controls (n=7) (12.56±2.70) vs. NMDA (n=38); p<0.001; Mann-Whitney). **h** Interevent interval of sEPSCs recorded from putative CA3 cells in slices prepared from animals after 7 days of NMDAR-ab (n=15 cells) as compared to the same measurements taken at 48 hours after NMDAR-Ab infusion (n=18 cells) (p=0.03; Mann-Whitney). **i** Amplitude measurements of sEPSCs recorded from putative CA3 cells in slices prepared from animals after 7 days of NMDAR-ab (n=15 cells) as compared to the same measurements taken at 48 hours after NMDAR-A infusion (n=18 cells) (p=0.03; Mann-Whitney). Measurements expressed as mean (M) ± standard error of the mean (SEM).

To confirm that these findings were concordant with the *in vitro* findings at 48h, we recorded LFPs in hippocampal brain slices at the completion of the recording experiments (ranging from day 7 to 14). In line with the spontaneous epileptiform activity seen *in vivo*, there was a considerable increase in the number of interictal events and a reduced IEI of ictal spikes in NMDAR-Ab infused animals compared to controls (Figure 3f,g). CA3 hippocampal whole-cell patch clamp experiments in NMDAR-Ab treated animals showed an even further increase in the IEI of sEPSCs (Figure 3h), and further reduction of the amplitude (Figure 3i) from that seen at 48 hours. Thus a reduction in excitatory neurotransmission was found to be coincident with the spontaneous epileptic events in both the 48 hour and 7 day NMDAR-Ab models.

### Reduced excitatory neurotransmission contributes to EEG changes

We tested whether the *in vitro* changes in single neuron behaviour associated with NMDAR-Ab contributed to *in vivo* (interictal) EEG patterns by using a computational model of microcircuit dynamics. We first generated quantitative predictions of the population-level effect of NMDAR-Ab based on *in vitro* differences between NMDAR-Ab and control-Ab exposed neurons measured using whole-cell patch clamp recordings (Figure 4a,b). We then fitted a four-population neural mass model (the canonical microcircuit (27)) to EEG power spectral densities, first for control animals, then for NMDAR-Ab animals under different empirical priors. These empirical priors were based on posteriors from the control-Ab treated animals and, for subsets of parameters, the quantitative predictions made from the *in vitro* recordings (see Materials and Methods). We then performed Bayesian model comparison of these models using a free energy approximation of the model evidence (Figure 4c). This comparison provides two key insights: (i) The model equipped with the full set of quantitative predictions from the microscale provides a more parsimonious explanation for the observed EEG spectral differences between control-Ab and NMDAR-Ab, compared to a null model without these microscale-derived predictions; and (ii) in a model space comparing models with only subsets of parameters informed by microscale-derived priors, the best model incorporates microscale-derived priors only for amplitude and population variance in one compartment of the canonical microcircuit model.

**Figure 4:**
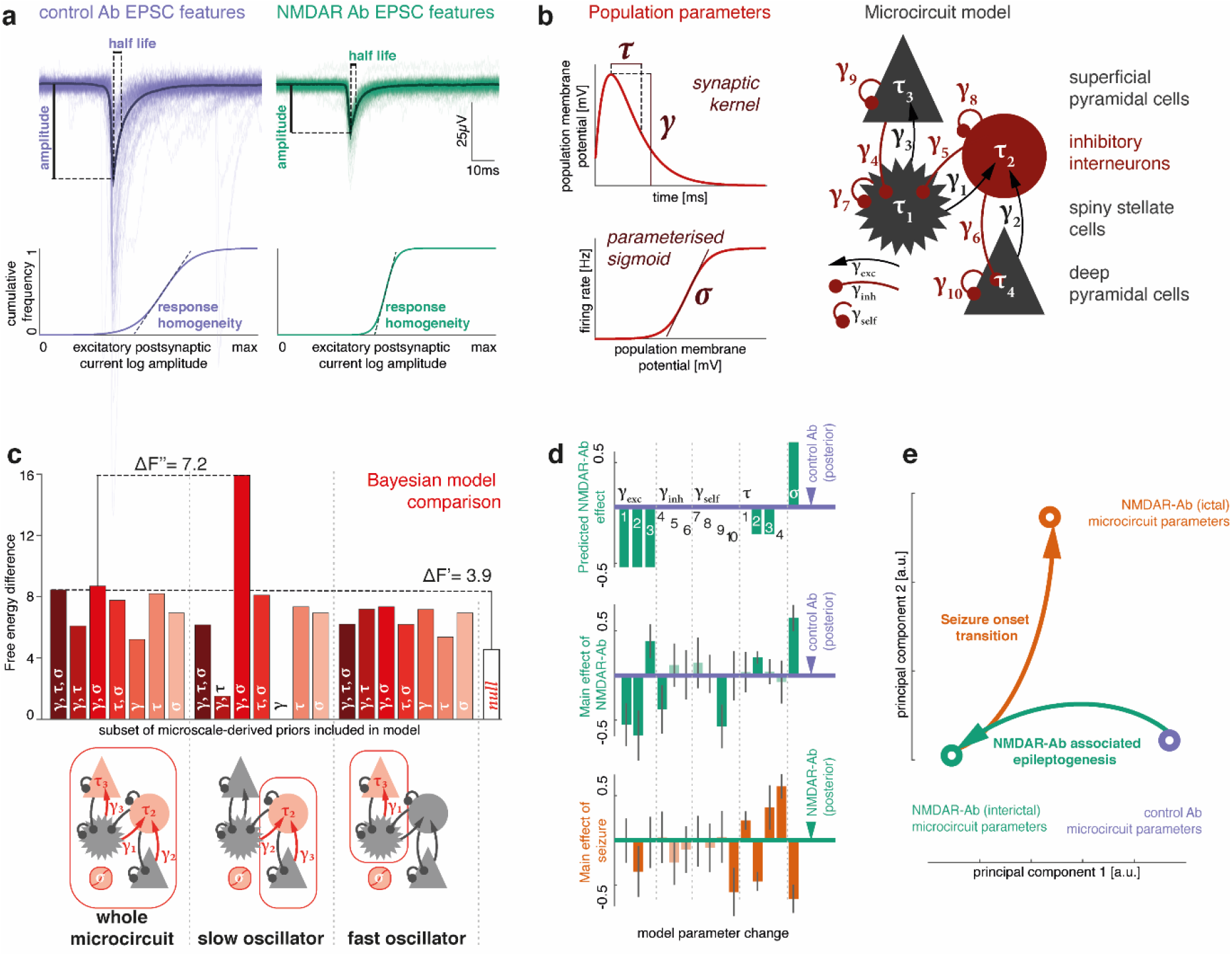
*In vitro* microscale disruptions of excitatory neurotransmission explain NMDAR-Ab-associated *in vivo* neurophysiological changes. **a** *In vitro* recordings of sEPSCs were used to quantify differences between control-Ab and NMDAR-Ab related neurotransmission, including half-life and amplitude of EPSCs. We also estimated EPSC homogeneity by fitting a sigmoid curve to the cumulative frequency of EPSC amplitudes, where the slope at the midpoint increases with more homogeneous EPSC amplitude distribution. **b** Microscale data features map onto population parameters in a canonical microcircuit model consisting of two layered neuronal oscillator pairs with recurrent excitation-inhibition coupling. Key synaptic parameters of this model are population synaptic response amplitude, time constant and response sigmoid, which are affected by EPSC amplitude, time constant, and amplitude distribution, respectively. Using the dynamic causal modelling framework, we fit a single such microcircuit to local field potential recordings in control Ab and NMDAR-Ab exposed mice. **c** Bayesian model comparison is shown for inverted models of NMDAR-Ab (interictal) local field potential dynamics where subsets of the excitatory synaptic parameters were informed by priors derived from microscale recordings. There is decisive evidence that the full model, where all parameters had microscale-informed priors, provides a more parsimonious explanation for the data than the model without predictive priors (BF ~ ΔF’ = 3.9). Of all tested models, the model containing predictive priors only for amplitude (γ) and population variance (σ), and only for the slow neuronal oscillator populations (inhibitory interneurons and deep pyramidal cells) performed best (BF ~ ΔF’ = 3.9) with a posterior probability of >95% in the given model space. **d** Parameter differences between conditions are shown for control vs. interictal NMDAR-Ab-associated dynamics (top: predicted; middle: estimated in the winning model), and seizure vs. interictal NMDAR-Ab dynamics (bottom: estimated). Error bars indicate a Bayesian 95% credible interval. Darker shading parameters where the estimated difference exceeds the credible interval. **e** A principal component decomposition of the full parameter set across three states (control-Ab, NMDAR-Ab (interictal), and NMDAR-Ab (seizure)) shows separability of the main effects of antibody (with most of the difference along the first principal component) and seizure onset (with most of the difference along the second principal component).

We then considered the overall winning model highlighted in Figure 4c and used its posterior parameters as priors for fitting EEG power spectral densities for seizure recordings. This modeling approach thus provided us with a parameter set capturing the difference between interictal EEG spectra in NMDAR-Ab and control-Ab treated animals (Figure 4d, middle panel), and between ictal EEG spectra and interictal EEG spectra in NMDAR-Ab (Figure 4d, bottom panel). Plotting these condition-specific parameterisations on a two-dimensional representation (Figure 4e) demonstrated separability of the two effects with near orthogonal effects of NMDAR-Ab (i.e. ‘epileptogenesis’) and seizure onset transition (i.e. ‘ictogenesis’).

### NMDAR-Abs push brain microcircuits into an unstable regime

To test whether there is a relationship between the changes induced by NMDAR-Ab and the increased propensity for spontaneous epileptic seizures in NMDAR-Ab infused animals, we ran simulations across an ‘epileptogenesis’ x ‘ictogenesis’ parameter space. We derived initial parameterisations by fitting microcircuit models to the three empirically observed brain states as described above (control-Ab, interictal NMDAR-Ab, and seizure NMDAR-Ab; model fits shown in Figure 5a). We then simulated linearly spaced intermediate steps between states along this 2-dimensional parameter space. At each point, we simulated the full spectrum, and classified it into one of the three empirically observed brain states by least mean squared difference between the simulated and observed spectra (Figure 5b). This map of parameter space shows that for microcircuits at control-Ab parameterization, the transition to a putative seizure-like territory is further away on the ‘ictogenesis’ parameter axis than it is for microcircuits at the interictal NMDAR-Ab parameterization. Each of the territories is characterised by particular spectral features, illustrated in Figure 5c by a map of high frequency and low frequency amplitude ratios across the same parameter space.

**Figure 5.**
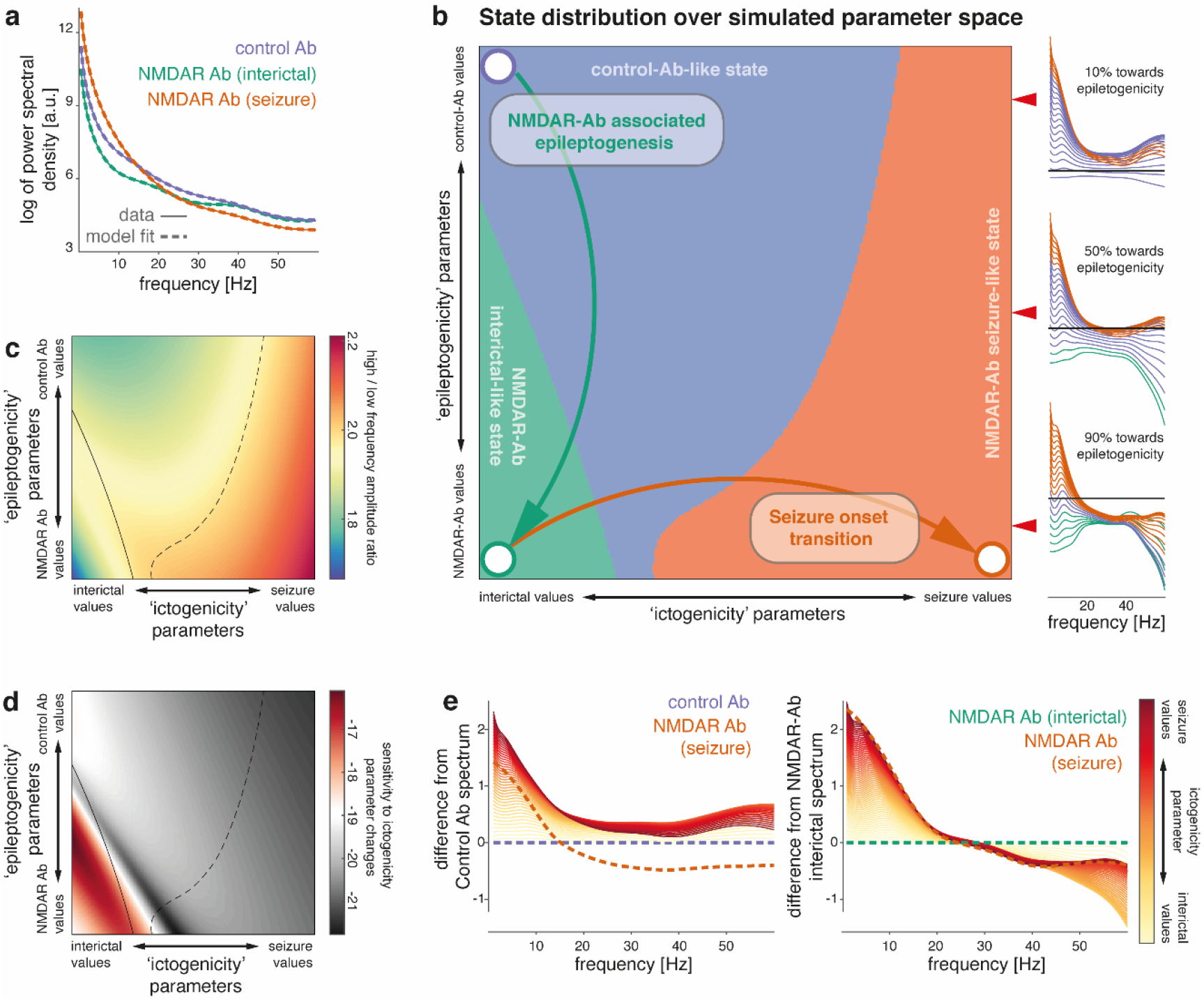
NMDAR-Ab pushes neuronal microcircuits towards unstable regimes. **a** Power spectra of LFP recordings of control-Ab injected mice, and interictal and ictal states of NMDAR-Ab injected mice are fitted with neural mass models of neuronal microcircuits. Dashed lines represent the model fit; solid lines represent the empirically measured spectra. **b** Based on the parameters derived from the dynamic causal model fits (represented here by circles), parameter changes that capture NDMAR-Ab associated alterations in microcircuitry (‘epileptogenicity’) and seizure onset transition (‘ictogenicity’) were identified (arrows). Linear combinations of these parameters are then used to simulate spectra at different locations in a two-dimensional parameter space along both ‘epileptogenicity’ on the y-axis, and ‘ictogenicity’ on the x-axis. At each point full spectra were generated, and classified as one of three possible states (‘Control-Ab-like’, ‘NMDAR-Ab interictal-like’, ‘NMDAR-Ab seizure-like’) based on the smallest mean-squared difference to the three empirically measured spectra. Insets show example simulations (normalized to the ‘Control-Ab’ spectrum, indicated by the black line) at three different values for ‘epileptogenicity’ indicated by the red arrow heads (10%, 50%, 90% towards full ‘epileptogenicity’ parameter change). Each line represents a simulation at different ictogenicity values spanning the full width of ictogenicity parameters, colour-coded by their classification. **c** Along the ‘epileptogenicity’ vs. ‘ictogenicity’ parameter space, the ratio of high frequency (>20Hz) and low-frequency (<8Hz) log amplitude is shown with state separating boundaries from panel (**b**) overlaid. **d** This plot demonstrates sensitivity to changes in ictogenicity parameters. For each given ‘epileptogenicity’ (i.e. row in the parameter space plot), mean-squared differences between neighbouring values of ‘ictogenicity’ are shown along the log-scale (a.u.) colour axis. State separating boundaries from panel (**b**) are overlaid. There is considerable overlap between regions in parameter space, where the microcircuit is sensitive to perturbation, and the NMDAR-Ab-like state identified in panel (**b**). **e** To demonstrate differences in sensitivity to ‘ictogenicity’ parameter changes, spectral differences from both control Ab parmeterisation and NMDAR Ab parameterisation are shown for increasing contribution of ‘ictogenicity’ parameters.

To evaluate sensitivity of these microcircuits to changes in the ictogenicity parameters, we quantified the difference of each simulated spectrum to spectra simulated from similar, ‘neighbouring’ ictogenicity parameter values (Figure 5d). This analysis revealed that the NMDAR-Ab interictal state is most sensitive to small changes in ictogenicity parameters.

Spectral changes associated with the increasing effect of ictogenicity parameters are further shown in Figure 5e, relative to control-Ab and interictal NMDAR-Ab microcircuit parameterisations.

### Pregnenolone sulphate (PregS) rescues NMDAR-Ab epileptic changes *in vitro*

We hypothesised that increasing NMDAR expression could rescue the electrophysiological features associated with NMDAR-Ab. The neurosteroid pregnenolone sulphate (PregS) has a membrane delimited action to increase the trafficking of functional NMDARs to the neuronal membrane (28). Using whole-cell patch clamp recordings of sEPSCs from untreated Wistar rat hippocampal slices we showed that PregS significantly increased amplitude and frequency of sEPSCs; the increased frequency was reversed by application of the NMDAR antagonist MK801 confirming NMDAR mediated effect (Supplementary Figure 2a-d). PregS reduced sIPSC IEI (162 ± 47.5 vs 101 ± 36.7, n=9; p=0.03, Wilcoxon paired rank test), but did not affect the amplitude (22.81 ± 2.5 vs 30.90 ± 3.2, n=9; p=0.25, Wilcoxon paired rank test) in control slices.

Next we tested whether these effects would rescue the NMDAR-Ab induced abnormalities observed *in vitro*. We recorded LFP discharges *in vitro* before and after addition of PregS to brain slices from animals treated with NMDAR-Ab or control-Ab for 48 hours or 7 days (Figure 6a). The number of interictal/spike events in NMDAR-Ab treated slices was significantly reduced after application of PregS and there was a corresponding increase in IEI (Figure 6b,c). By contrast, slices from rats treated with control-Ab showed no significant reduction in interictal events after application of PregS, although the IEI did show a measurable reduction post PregS (Figure 6b,c).

**Figure 6.**
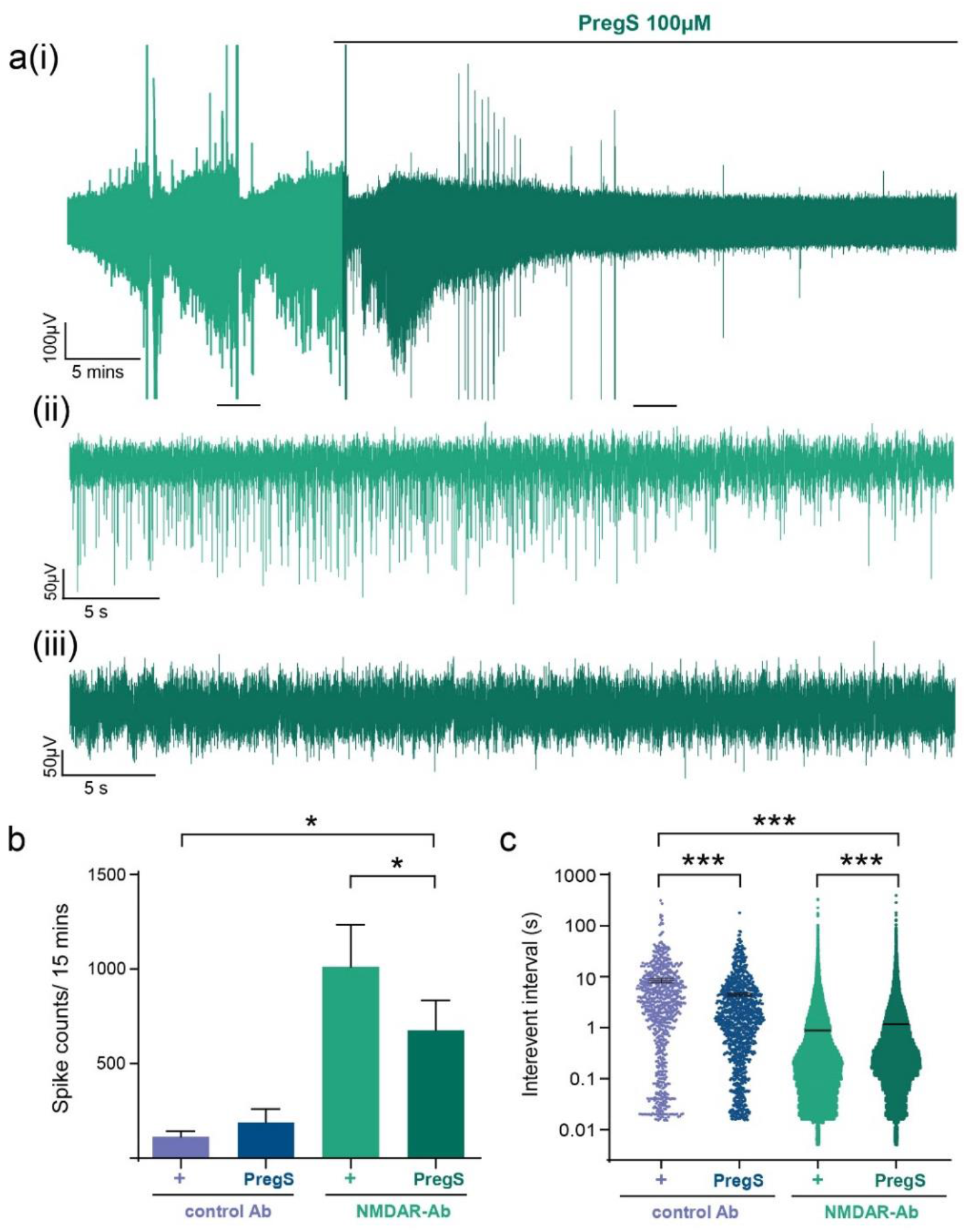
Pregnenolone sulphate reduces NMDAR-Ab mediated spontaneous ictal activity *in vitro*. **a** Representative *in vitro* slice LFP recording (subpanel **a**i) from a 7-day NMDAR-Ab treated animal illustrates the response to PregS 100μM. Scale bar 10μV vs. 5 mins. Subpanels **a**(ii) and **a**(iii) represent the magnification of control and PregS 100μM trace area respectively. Scale bars 50μV vs. 5 s. **b** Ictal spike counts after PregS is applied to control-Ab infused slices (n=6 slices; p=0.07, Wilcoxon paired rank test) and NMDAR-Ab infused slices (n=45 slices; p<0.05, Wilcoxon paired rank test). **c** IEI recorded in the control Ab infused LFP slices after addition of PregS indicates an increased frequency of events (n=6 slices; p<0.001, Wilcoxon paired rank test). The frequency of events is reduced in the NMDAR-Ab slices after addition of PregS (n=36 slices; p<0.001; Wilcoxon paired rank test). Measurements expressed as mean (M) ± standard error of the mean (SEM).

We confirmed that the reduction in LFP spiking activity was associated with a rescue of the synaptic abnormalities by performing whole-cell patch clamp recordings in hippocampal slices from animals exposed to short (48 hours) or more chronic (7 days) NMDAR-Ab exposure. In both conditions, PregS increased the frequency of sEPSCS recorded in CA3 neurons from NMDAR-Ab treated animals back to control levels (as measured by a fall in IEI; Figure 7a,b). There was no significant effect of PregS on sEPSC amplitude (15.2 ± 0.93 vs 14.5 ± 1.3, n=12; p=0.5, Wilcoxon ranked pairs test), mini-IPSC amplitude (18.9 ± 4.1 vs 15.9 ± 2.6pA, n=6 in both groups; p=0.15, Wilcoxon ranked pairs test) or mini IPSC IEI (200 ± 39 vs 60.2 ± 69.3 ms, n=6 in both groups; p=0.22, Wilcoxon ranked pairs test).

**Figure 7.**
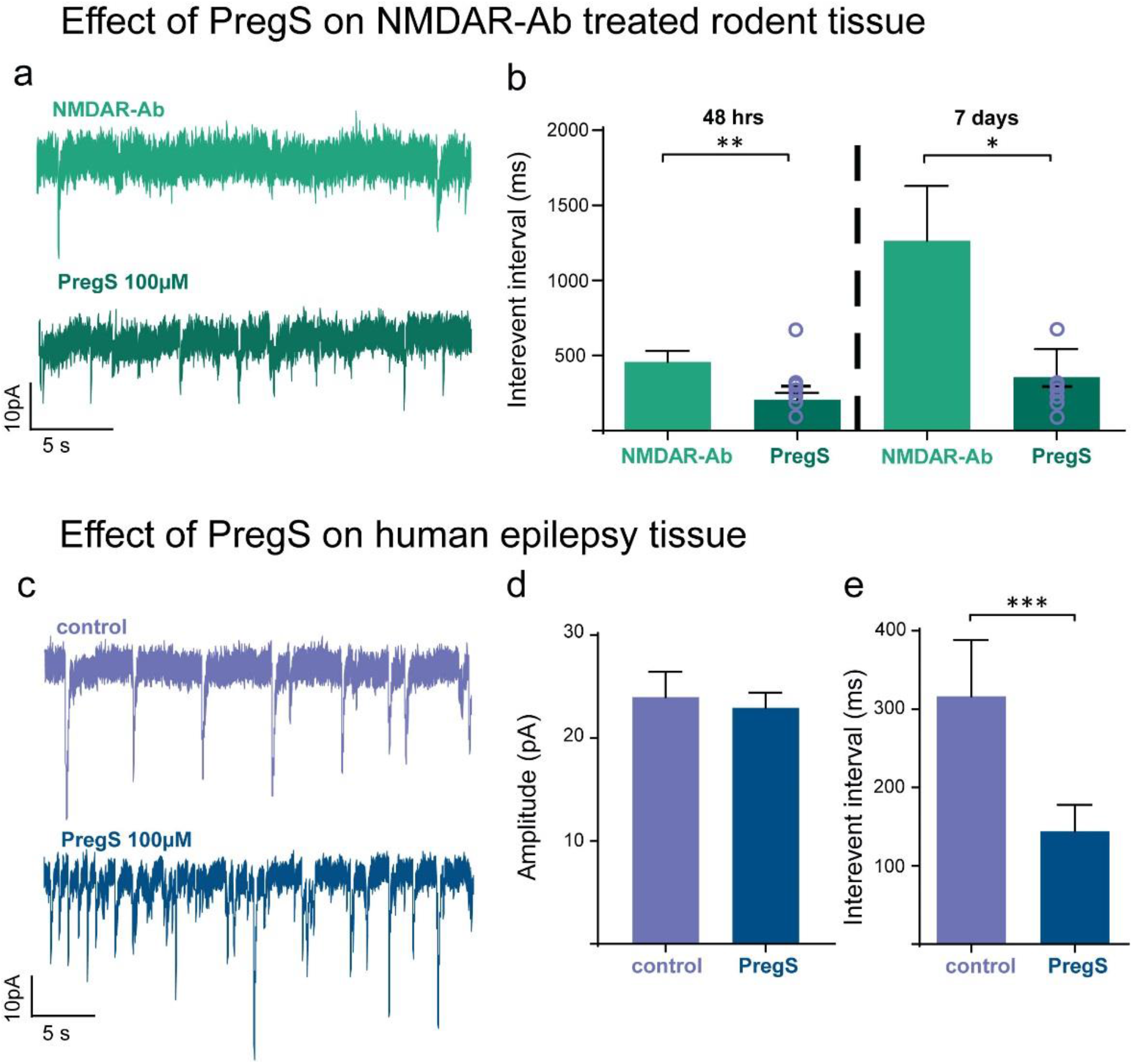
Pregnenolone sulphate increases glutamatergic transmission in NMDAR-Ab infused rodent hippocampal slices *in vitro* and has similar effects in human epilepsy brain tissue. **a** Representative whole-cell patch clamp recording of putative CA3 pyramidal cell sEPSCs from an NMDAR-Ab treated hippocampal slice before and after addition of PregS. Scale bar 10pA vs. 5s. **b** IEI of sEPSCS recorded from putative CA3 pyramidal cells before and after PregS application to hippocampal slices prepared from NMDAR-Ab treated animals at 48 hours after Ab injection (n=12 pairs; p=0.0093, Wilcoxon paired rank test) and after 7 day NMDAR-Ab infusion (n=6 pairs; p=0.03, Wilcoxon paired rank test). The IEI of sEPSCs recorded from CA3 pyramidal cells in control-Ab slices are added for comparison (n=7; shown as open grey circles superimposed on PregS bar) at the 48 hour (p=0.55, Mann-Whitney test) and 7 day timepoint (p=0.8, Mann-Whitney test). **c** Representative *in vitro* whole-cell patch clamp recordings of sEPSCs in human paediatric epilepsy before and after addition of PregS. Scale bar 10pA vs. 5s. **d** Amplitudes of sEPSCs recorded from cells in human brain tissue before and after addition of PregS *in vitro* (n=9; p=ns (1), Wilcoxon ranked pairs test). **e** Interevent interval of sEPSCS recorded from cells in human brain tissue before and after addition of PregS *in vitro* (n=9; p=0.004, Wilcoxon ranked pairs test). Measurements expressed as mean median (M) ± standard error of the mean (SEM).

Finally, we showed that this acute reversal of NMDAR-Ab mediated electrophysiological effects by PregS could in principle be translated to human brains. Pediatric brain slices prepared from tissue resected during epilepsy surgery (Figure 7c) was exposed to PregS. As in the rat brain slices, we saw a similar increase in the frequency of sEPSCs (i.e. reduction of IEI of sEPSCs) but no effect on the sEPSC amplitude (Figure 7d,e). There was also no significant change in the sIPSCS amplitude after addition of PregS to human tissue (31.87±9.7 vs 27.96±4.1, n=6; p=0.44, Wilcoxon paired rank test), or in the IEI (123 ± 41.8 vs 95.7 ± 15.8; p=0.09, Wilcoxon paired rank test).

## Discussion

NMDAR-Abs disrupt normal brain circuit function and are associated with a range of neurological symptoms in patients, including epileptic seizures. The majority of these symptoms are now understood to be directly caused by antibody-induced hypofunction of the excitatory NMDAR. However, the link between reduced NMDAR function and increased seizure propensity has remained unexplained, and is particularly counterintuitive from a classical perspective of seizures arising from an excitatory-inhibitory imbalance (29). Combining electrophysiological recordings in a rat model of NMDAR-Ab encephalitis and computational modeling of brain microcircuit function, we show that an effective reduction in excitatory synaptic neurotransmission observed at the single-neuron scale underlies NMDAR-Ab epileptogenesis by altering the dynamical makeup of brain microcircuitry, making it more susceptible to excursions into pathological brain states. The modelling work here illustrates a clear separation between ictogenesis, and epileptogenesis. The transition from interictal to ictal state (i.e. ictogenesis) may indeed be best explained by changes in synaptic coupling that effectively push excitation-inhibition out of balance into a hyperexcitable state. Yet the pathophysiological process of epileptogenesis – i.e. the synaptic changes that render the brain more likely to undergo an interictal-ictal transition – are separable from those associated with ictogenesis. In fact, in our particular animal model we can show that the *hypo*excitation induced by NMDAR-Ab counterintuitively contributes to epileptogenicity, by increasing the sensitivity of microcircuits to fluctuations of the ictogenicity parameters. We demonstrate that this phenotype can be rescued with the neurosteroid PregS, an agent known to increase NMDAR availability at the neuronal membrane. Furthermore, we reproduced the synaptic effect of PregS in human brain tissue *in vitro*, indicating its potential use as precision-treatment for epileptic seizures in patients with established NMDAR-Ab encephalitis and potentially other seizure aetiologies associated with NMDAR-hypofunction.

### Modelling NMDAR-Ab epileptogenesis in rats

The genetic background of rodent models can confer different levels of seizure susceptibility (30, 31). Juvenile Wistar rats are highly seizure susceptible at P21 (32) and proved to be an effective choice for this passive transfer NMDAR-Ab-mediated seizure model. The CA3 region was found to have the most spontaneous epileptic activity after NMDAR-Ab ICV injection, and epileptic discharges were therefore recorded from this location *in vivo*. The CA3 hippocampal region is also the seizure-onset zone in other murine models of epilepsy (33, 34). Another study using NMDAR-Ab positive CSF recorded a higher incidence of CA3 epileptiform afterpotentials in brain slices from Wistar rats injected with NMDAR-Abs compared to controls (35); these epileptiform events were not seen in other hippocampal areas (36, 37). Furthermore, electrophysiology studies with Leucine-rich-glioma-1 autoantibodies (LGI1-abs), that cause facio-brachial dystonic seziures and limbic encephalitis enhanced CA3 pyramidal cell excitability when applied *in vitro* (38, 39), and patients show focal CA3 subfield atrophy on 7T MRI following LGI1-ab mediated disease (40). Together, these reports and our results confirm that the CA3 subfield is an important epileptogenic hippocampal region in immune-mediated types of epilepsy, justifying its further investigation.

### Reductions in excitation can increase seizure propensity

Almost all healthy brains can produce epileptic seizures in the context of acute excitation-inhibition imbalance, such as acute GABA-blockade caused by chemoconvulsants (41). Furthermore, many effective anti-seizure drugs aim to restore this balance by increasing levels of inhibition or reducing neuronal excitation (42). Hypofunction in both excitatory AMPAR and NMDA receptors have been reported in epilepsies and seizure models, including antibody-induced loss of AMPARs (43, 44). Similarly, recent work in a rat model of chronic temporal lobe epilepsy (TLE) demonstrates that dysfunctional regulation and resultant hypofunction of GluA1 (an AMPAR subunit) is an important factor in both epileptogenesis and the spread of seizures across the brain (45). Defects of another AMPAR subunit, GluA2, are also found in neurodevelopmental disorders such as epileptic encephalopathy in children (46).

Genetically modified mice lacking NMDARs in CA3 pyramidal neurons were more susceptible to kainate-induced seizures, and pharmacological blockade of CA3 NMDAR in adult wild-type mice produced similar results (47). Loss of function mutations in both the GluN1 and GluN2A subunits of the NMDAR have been identified in patients with epilepsy and neurodevelopmental disorders (48, 49). A recent dynamic causal modelling study of NMDAR-Ab encephalitis patient EEGs also showed that the deficit in signaling at NMDARs in patients with NMDAR-Ab encephalitis was predominantly located at the NMDA receptors of *excitatory* neurons rather than at inhibitory interneurons (50), as shown here in our model. By combining detailed multiscale electrophysiology and computational modelling, we provide further evidence that the direct synaptic effects of NMDAR-Ab identified from whole-cell patch clamp recordings offer a quantitative explanation for the epileptic brain dynamics captured *in vivo* using local field potentials.

In our circuit models of NMDAR-Ab-induced epileptogenesis, the parameter changes required to optimally model the associated LFP changes required not only changes in NMDAR-transmission related parameters, but also more widespread changes involving both excitatory and inhibitory neuronal populations. Two possible explanations for this, among others, include heterogeneous expression of the molecular deficit and emergent properties in a complex system. Indeed compensatory and maladaptive network changes could also contribute to the increase in pyramidal neuron and circuit excitability through increased activity at other glutamatergic receptors (i.e. AMPA and kainate receptors), as observed in some mouse models (51). Regardless of the mechanisms, the subacute evolution of seizures suggests that plasticity induced by the reduction in NMDAR-Ab transmission plays an active role in epileptogenesis.

Heterogeneity of NMDAR distribution at normal baseline function may also account for the unexpected effect of NMDAR-hypofunction in neuronal circuit dynamics. Selective forebrain pyramidal neuron NR1 knockout mice demonstrated both socio-cognitive deficits and *increased* pyramidal cell excitability (52), potentially driven by a release from inhibition by reducing putative NMDAR-mediated excitatory driving of inhibitory neuron activity, or by changing phase relationships between interacting neuronal populations. Similarly, hippocampal CA3 NMDA receptors may normally act to suppress the excitability of the recurrent network by restricting synchronous firing of CA3 neurons (47), with NMDAR-blockade releasing them from this suppression. Our modelling also suggests that NMDAR-Ab hypofunction may have stronger impacts on specific compartments of the affected microcircuitry.

Lastly, the integrated brain (micro-)circuits under investigation here show complex emergent dynamics arising from a range of often nonlinear interactions. This complexity can often give rise to unpredictable results, even where simple systems show a clear mapping between excitation/inhibition balance and seizure propensity. Our modelling suggests that the effective synaptic connectivity changes associated with NMDAR-Abs sensitise the circuit to perturbations in another (largely unrelated) set of parameters. This insight implies that whilst the seizure transition may be indicative of a change in a balance of parameters associated with excitation and inhibition, there may be territories of parameter space where this change is tolerated without any notable impact on the local field potential and other territories where the same change of excitation/inhibition parameter balance would produce large changes consistent with an epileptic seizure. The simulations here indicate that the NMDAR-Ab models are much more sensitive to ‘ictogenesis’-associated parameter changes, suggesting that the combined parameter changes induced by NMDAR-Ab have pushed the microcircuit into a territory in parameter space, where it is particularly sensitive to further perturbations.

### Neurosteroid treatments

Neurosteroids are derived from cholesterol and are precursors to gonadal steroid hormones and the adrenal corticosteroids. They are produced *de novo* by glial cells and principal neurons, hence the term ‘neurosteroid’ (53), and modulate brain excitability primarily by interacting with receptors and ion channels on the membrane surface rather than within cells (54). Pregnenolone (precursor to PregS) elevates levels of allopregnanolone and pregnenolone sulphate, a positive NMDAR receptor modulator. Pregnenolone has been used as an effective adjunctive therapy in treatment trials for schizophrenia, a psychiatric condition that shares the pathophysiology of “NMDAR hypofunction” (55–57). These trials demonstrate both the safety and efficacy of Pregnenolone in humans. *In vivo*, pregnenolone was found to reduce the hyperlocomotion, stereotypic bouts and prepulse inhibition deficits in a mouse model of schizophrenia acutely, and improved impaired cognitive deficits with chronic treatment (58). PregS, a sulfated ester of pregnenolone, is a positive NMDAR modulator (59, 60) and studies show that the mechanism of action is to increase surface expression of functional NMDARs containing the NMDAR subunit subtype 2A (NR2A) or subtype 2B (NR2B), both the most abundant subunits found in the brain; no evidence points to changes in NR2C or NR2D subunits (28). In a previous study, this membrane delimited effect postulated to be via a non-canonical, G-protein and calcium-dependent mechanism occurred within ten minutes of application to cortical neurons, increasing surface levels by 60-100% (28); this speed of action of PregS was reproducible *in vitro* in our brain slice recordings. In addition to the effect of PregS on NMDARs, other CNS targets have been described, including the GABA_A_ receptor, glycine receptor, and more recently the TRPM3 channel (61) and Kir2.3 channel (62). The effects of PregS on the GABAAR most commonly described are at lower, nanomolar range doses (63, 64). We found that the 100μM dose of PregS had no effect on GABAergic neurotransmission in the brain slices from the NMDAR-Ab model. A proconvulsant effect was seen in the control antibody slices, but the reverse was seen in the NMDAR-Ab infused brain slices, suggesting that it was the increase of NMDARs that led to the reduction in the frequency of epileptic events.

Previous studies with NMDAR modulators and other treatments have focused on the prevention of pathogenic antibody effects (9–11). To our knowledge this is the first study to show that symptoms established for more than 48 hours (in this case ictal activity *in vitro*), can be treated with the use of the neurosteroid PregS. Whilst PregS was chosen here specifically to treat the known underlying pathophysiology, its role for the treatment of epilepsy may extend into other conditions or scenarios where the treatments that accord to the notion of reestablishing excitation-inhibition balance have failed to be effective (e.g. refractory status epilepticus) (65).

### Limitations

We did not record directly from inhibitory interneurons but it is likely that both cell types are involved in driving the epileptogenesis. The CA3 network is highly complex, with CA3 pyramidal cells having convergent and divergent connections with several interneuronal types in different hippocampal regions (66). A single CA3 pyramidal cell can make synapses with 30,000-60,000 neurons in the ipsilateral hippocampus (67). It therefore remains a possibility that NMDAR hypofunction affecting GABAergic interneuronal populations produces a disinhibited CA3 network (52, 68, 69) and investigating this possibility should be a goal of further electrophysiological studies.

Another limitation to this study was that the cognitive effects of NMDAR-Abs were not investigated in the animals or electrophysiologically through long-term potentiation (LTP) recordings. However, LTP recordings will be difficult to interpret with spontaneous ictal spikes as seen in these brain slices. It is known that in CA3, NMDAR-dependent (associational-commissural, A-C fibres) and NMDAR-independent (Mossy-fibre) forms of LTP are expressed in adjacent synapses. LTP has been shown to be reduced by NMDAR-Ab positive CSF (experiments performed in Wistar rats), and this reduction may contribute to the cognitive deficits seen clinically (35). Mossy fibre synapses display a markedly lower proportion of GluN2B-containing NMDARs (70) than associative or commissural synapses, therefore a loss of NMDARs in this critical population of cells could also be devastating. PregS offers an attractive treatment strategy to increase NR1 and NR2B subunit NMDARs (28) that could potentially mitigate the short and long-term cognitive effects seen in patients, as well as the seizures. Further detailed molecular studies will need to be done to characterize the NMDA receptor and subunit changes.

### Outlook

Future studies need to focus on ascertaining the optimal brain concentrations of the neurosteroid PregS *in vivo* in the NMDAR-Ab mediated disease models, and on determining whether these can be achieved through direct administration of PregS (e.g., by subcutaneous injection) or the precursor Pregnenolone (oral administration, a treatment already used safely in humans)(57). There is precedence for the use of neurosteroids in other forms of epilepsy, including genetic epilepsies (e.g., Phase 3 clinical trial of ganaxolone in PCDH-19 female patients; NCT03865732) as well as refractory status epilepticus (71). In genetic epilepsy, research efforts have focused for some time on modifying the specific underlying genetic effect with new and repurposed drugs (72). In this study, we show that there is also a possibility of providing receptor-specific neurosteroid treatment to affected patents with immune-mediated epilepsy and encephalitis.

## Materials and Methods

### In vitro experiments

#### NMDAR-Ab and control antibodies

Three human-derived NMDAR-Ab preparations were obtained for experimental use. The first was NMDAR-Ab positive IgG prepared from plasma obtained, with informed consent at plasmapheresis from a patient (11 year old female with NMDAR-Ab encephalitis), using Protein G beads as previously described (18). In addition, two different recombinant NMDAR-Ab specific human monoclonal antibodies were used; SSM5 (preparation detailed in (8)) and 003-102 (preparation detailed in (3)). Both of these NMDAR-Ab monoclonal antibodies have previously been shown to have *in vitro* and *in vivo* pathogenic effects (3, 8).

Control antibodies (Control-Abs) were human derived IgG from one healthy individual (healthy male, aged 35 years), and two non-brain reactive monoclonal Control-Abs; mGo53 (3), and 12D7 (8).

As all antibody preparations had previously been used in the studies specified and were known to be pathogenic, and stocks for some preparations were limited, a direct comparison was not made between each antibody preparation of the epileptogenic effects. The three NMDAR-Ab preparations will be collectively referred to as “NMDAR-Abs” throughout the paper and the three control preparations as “Control-Abs”. Detailed description of antibody use per experiment is provided in Supplementary Table 1.

#### Animals

Given our experience in other rodent epilepsy models (32, 45), we hypothesized that the use of juvenile male Wistar rats would increase the probability of observing spontaneous seizures in a NMDAR-Ab passive transfer model. Thirty-one post-natal day 21 (P21) male Wistar rats, weighing 50-58g, were used for the *in vitro* experiments and 12 were used for the *in vivo* experiments. The animals were housed in temperature- and humidity-controlled conditions with a 12h/12h light/dark cycle, and allowed to feed and drink *ad libitum*. All procedures were compliant with current UK Home Office guidelines and ARRIVE guidelines.

#### Surgery: stereotaxic ICV injection of NMDAR- and control antibodies

ICV injection was performed as previously described (18). The co-ordinates for the left lateral ventricle in this model were 1.5mm left lateral and 0.6mm caudal from bregma (73). 10 micrograms (10mcg) of SSM5 (NMDAR-Ab monoclonal antibody), or non-reactive control antibody (12D7) or 8uL of NMDAR-Ab IgG (concentration 37mg/ml) (18) or healthy control IgG (concentration 30mg/ml) was used.

#### Local field potential (LFP) recordings

Forty eight hours after ICV injection or at the end of the recording period in telemetry experiments, rats were anaesthetized using isoflurane followed, after loss of consciousness, by pentobarbital (60 mg/kg, SC) and xylazine (10 mg/kg, IM). Transcardial perfusion was then performed using ice-cold artificial cerebrospinal fluid (cutting aCSF) in which NaCl had been replaced with equimolar sucrose. Animals were decapitated, and the brain was removed and placed in ice-cold cutting aCSF. Sagittal slices at 450μm were cut using a vibratome (Campden Instruments, UK) as described previously (74). The slices were transferred to a storage chamber containing standard aCSF (in mM: 125 NaCl, 3KCl, 1.6 MgSO_4_, 1.25 NaH_2_PO_4_, 26 NaHCO_3_, 2 CaCl_2_, 10 glucose) and constantly bubbled with carbogen (95% O_2_/5% CO_2_) at roomtemperature for at least one hour prior to recording. Slices were then transferred to an interface chamber (Scientific System Design Inc., Canada) and continuously perfused with aCSF (2-3ml/min) maintained at 30-31°C and visualised with a SZ51 stereomicroscope (Olympus). A Flaming-Brown micropipette puller (Sutter Instruments, CA) was used to pull borosilicate glass microelectrodes with an open tip resistance of 1-3 MΩ when filled with aCSF. Electrodes were inserted into CA3 and CA1 regions of the hippocampus using Narishige MM-3 (Narishige, Japan) micromanipulators and signals were recorded using an EXT-02 headstage and amplifier (NPI Electronic). HumBugs (Quest Scientific) were used to remove electrical 50 Hz interference from recordings. The signal was then amplified x100 via an EX10-2F amplifier (NPI Electronics) and filtered (700 Hz low-pass, 0.3 Hz high-pass, sampling rate 5 kHz) and further amplified x10 using a LHBF-48X amplifier (NPI Electronics). Signals were then digitised at 10 KHz using an analogue to digital converter (Micro-1401-II; CED). LFP recordings were assessed using Spike2 software (CED) for spontaneous epileptiform activity. Spike2 was used to calculate the root mean square (RMS) amplitude of each recording. Epileptiform activity was classified as an event when it displayed an amplitude greater than four-fold the RMS amplitude, providing the event count, while the time difference between these events provided the interevent interval. A custom MATLAB script was used to prevent false event detection, removing those with an interevent interval less than 15ms. Statistical analysis was conducted using Graphpad Prism 8 (GraphPad Software Inc). Measurements were expressed as mean (M) ± standard error of the mean (SEM).

#### Whole-cell patch clamp recordings

For whole-cell patch clamp recordings, brain slices were prepared as above but sliced at a thickness of 350μm. For recording, the slices were transferred to a submerged chamber (Scientifica, UK) and visualised using Nomarski infra-red optics on a BX51WI Microscope (Olympus). A Flaming-Brown micropipette puller (Sutter Instruments, USA) was used to pull borosilicate glass microelectrodes of 4-6 MΩ for recordings. Electrodes were filled with an internal solution containing (in mM): 100 CsCl, 40 HEPES, 1 Qx-314, 0.6 EGTA, 5 MgCl_2_, 10 TEA-Cl, 4 ATP-Na, 0.3 GTP-Na and 1 IEM 1460 (titrated with CsOH to pH 7.25) at 290-295 mOsm for IPSCs. For EPSCs the internal solution contained (in mM): 40 HEPES, 1 Qx-314, 0.6 EGTA, 2 NaCl, 5 Mg-gluconate, 5 TEA-Cl, 10 Phospho-Creatinine, 4 ATP-Na, 0.3 GTP-Na (titrated with CsOH to pH 7.3) at 285 mOsm for EPSCs. The EPSCs and IPSCs were recorded as apparent inward currents at −70 mV using an Axopatch 200B amplifier (Molecular Devices, USA). Signals were low-pass filtered at 5kHz with an 8-pole Bessel filter and digitised at 10kHz using a Digidata 1440A and pClamp software (Molecular Devices). Data were analysed using Axograph and Prism 8. Measurements are expressed as mean median ± SEM.

For human tissue recordings, brain slices were prepared as previously described (75, 76). Briefly, human tissue was obtained with informed parental or guardian consent from pediatric patients undergoing epilepsy surgery at Birmingham Children’s Hospital. Ethical approval was obtained from the Black Country Local Ethics Committee (10/H1202/23; 30 April 2010), and from Aston University’s ethics committee (Project 308 cellular studies in epilepsy) and through the Research and Development Department at Birmingham Children’s Hospital (IRAS ID12287). Brain tissue was obtained from 7 patients (F:M 4:3), median age 6 years (range 4-16 years). Surgical procedures included frontal resection (three), temporal parietal occipital disconnection (one), temporal lobectomy (two), and right hemispherectomy (one) as described in Supplementary Table 2. Specimens were resected intraoperatively with minimal traumatic tissue damage, and minimal use of electrocautery. Brain tissue was transported in ice-cold choline-based aCSF as previously described (76).

#### Drugs

For all *in vitro* electrophysiology experiments, Pregnenolone sulphate (Sigma, UK) was prepared as 1M stock using dimethyl sulfoxide (DMSO). A concentration of 100μM was used for all electrophysiology experiments. MK-801 (Hello Bio, UK) was prepared with distilled water as 10mM stock and used at 10μM concentrations.

#### Immunofluorescence and image analysis

Following *in vitro* electrophysiology experiments, brain slices were briefly fixed in 4% paraformaldehyde (PFA) for 45 minutes. To determine NMDAR-Ab or control antibody binding, the sections were rinsed with phosphate buffered saline (PBS) and then incubated in antihuman IgG Alexa-Fluor-488 at 1 in 1000 (Invitrogen, UK) overnight at 4·C. Sections were washed and mounted with aqueous mounting medium containing DAPI. Images of hippocampal sections were taken on a Tandem Confocal Scanning SP5 II microscope (Leica Microsystems Ltd) using a (10x/0.30) dry objective lens. Fluorescence was excited with a 488nm argon laser at 23% power (emission bandwidth 504nm-564nm). Confocal micrographs were acquired at 1024 pixels2 with actual area of 1.48mm2, and scanning speed 100 Hz.

Biocytin (Sigma, UK) was added to the electrode filling solution at a concentration of 5mg / ml to determine the characteristics of the recorded cell during whole-cell patch clamp experiments. Neurons were filled with biocytin and slices were fixed overnight at 4°C in 100 mM PBS (pH 7.3) containing 4% PFA (BDH, USA). After washing in PBS, slices were incubated for 18 hours in PBS supplemented with 1% Triton X-100 and 0.2% streptavidin Alexa Fluor 488 conjugate (Invitrogen) at 4-6°C. Sections were washed and mounted with aqueous mounting medium containing DAPI for imaging.

Fluorescent intensity Log EC_50_ ratio of hippocampal sections was determined using 5 regions of interest (ROI) of the same size from the molecular outer layer and from the granular cell layer of CA3. The mean fluorescent intensity Log EC_50_ value of the outer molecular layer was normalised by dividing it by the mean fluorescent intensity Log EC_50_ of nonspecific staining in the granular cell layer. This process provided a ratio for each replicate. All images were analysed through FIJI by converting them to grayscale (8-bit) before generating cumulative pixel intensity histograms for each ROI which were calculated using a customized macro programme (1, 18). Any statistical analysis performed was conducted in GraphPad Prism 8.

### In vivo experiments

#### Surgery: placement of ventricular catheters, osmotic pumps and wireless EEG transmitters

Osmotic pumps (model 1007D, Azlet) were used for cerebroventricular infusion of monoclonal antibodies and human IgG (volume 100μl, flow rate 0.5μl, duration 7 days) (7, 8). The day before surgery, two osmotic pumps per animal were prepared by loading with monoclonal antibodies or human IgG. The loaded pumps were then connected to polyethylene tubing 69mm-x-1.14mm diameter (C312VT; PlasticsOne) and a double osmotic pump connector intraventricular cannula (328OPD-3.0/SPC; PlasticsOne). Pumps were left overnight in sterile saline solution at 37°C. The next day, under isoflurane anaesthesia, rats were placed in a stereotaxic frame for surgery. The osmotic pumps were placed subcutaneously and an attached cannula was inserted into the lateral ventricles (1.5mm lateral, 0.6mm caudal). A subcutaneous pocket was formed over the right flank with a single skin incision and blunt tissue dissection for the transmitter (A3028B-DD subcutaneous transmitters, 90-mm leads, OpenSource Instruments (OSI)), and depth electrode (W-Electrode (SCE-W), OSI) placed in the left hippocampus (CA3, 3.5mm lateral, 3.6mm caudal, depth 2.3mm) with a reference electrode implanted in the contralateral skull (3.5mm lateral, 3.6mm caudal). The cannula and skull electrodes were secured with dental cement as previously described (18, 25).

#### Collection and analysis of EEG data

A custom-built Faraday cage with an aerial was used to collect and record EEG data. Transmitter signals were continuously recorded in animals while freely moving using Neuroarchiver software (OSI) and analysed as previously described (18, 25, 77). In brief, for automated ictal event detection, video-EEG matching was used to identify ictal EEG events. The Event Classifier (OSI) was then used to classify one second segments of EEG according to program metrics (power, coastline, intermittency, coherence, asymmetry, spikiness) enabling similar events to cluster together when plotted. This generated a library of ictal events that allowed fast identification of abnormal EEG events by automated comparison to the library (http://www.opensourceinstruments.com/Electronics/A3018/Seizure_Detection.html).

Powerband analysis was carried out using a custom-designed macro. Statistical analysis was conducted using Graphpad Prism 8 (GraphPad Software Inc.).

### Dynamic causal modelling

Dynamic causal modelling (DCM) was performed using the academic software package SPM12 (https://www.fil.ion.ucl.ac.uk/spm/). All analysis code and raw data are available online at http://doi.org/2010.17605/OSF.IO/GUPBF. Analogous to previous work(21, 78), we modelled LFP spectra recorded in rat hippocampal tissue as arising from a single canonical microcircuit at steady state (79). This ‘canonical microcircuit’ is a layered set of recurrently coupled excitatory and inhibitory neural masses (80–82). Originally conceived as a description of layered cortical population activity, the canonical microcircuit model has also been shown to offer a parsimonious explanation of rodent hippocampal activity in different physiological and abnormal states (83). The four populations of the canonical microcircuit act as two coupled oscillator pairs: One superficial oscillator typically characterised by fast (beta/gamma-range) frequencies and consisting of ‘superficial pyramidal cells’ and ‘spiny stellate cells’; and one deep oscillator with typically slow (theta/alpha-range) frequencies consisting of inhibitory interneurons and deep pyramidal cells. This model is a description of neuronal interactions at the population level. Effective synaptic connectivity is parameterized by three key parameters: effective coupling strength (γ), a postsynaptic time constant (τ), and the slope of a parameterised sigmoid function (σ) that encodes population response variance (84). The free neuronal parameters in the model implemented here are summarized in Table 1.

**Table 1:** Free parameters fitted by the DCM. (parameters related to NMDAR-transmission in bold)

|  |  |
| --- | --- |
| $\tau_1$ (spiny stellate time constant) | $\gamma_1$ (**ss to ii excitation**) |
| **$\tau_2$ (inhibitory interneuron time constant)** | **$\gamma_2$ (dp to ii excitation)** |
| **$\tau_3$ (superficial pyramidal cell time constant)** | **$\gamma_3$ (ss to sp excitation)** |
| $\tau_4$ (deep pyramidal cell time constant) | $\gamma_4$ (sp to ss inhibition) |
| **$\sigma$ (population response variance)** | $\gamma_5$ (ii to ss inhibition) |
| | $\gamma_6$ (ii to ip inhibition) |
| | $\gamma_7$ (sp inhibitory self-modulation) |
| | $\gamma_8$ (ii inhibitory self-modulation) |
| | $\gamma_9$ (ss inhibitory self-modulation) |
| | $\gamma_{10}$ (dp inhibitory self-modulation) |

DCM employs a standard variational Laplace model inversion to (i) estimate posterior densities of model parameters given the data and some prior values, and (ii) provide an estimate for the Bayesian model evidence using the free energy approximation (85, 86). This estimation of the Bayesian model evidence allowed us to compare evidence for models of the same data under different assumptions encoded in different priors. Specifically, we wished to test the hypothesis that NMDAR-Ab-induced changes measured using whole-cell patch clamp recordings contribute to the LFP features observed *in vivo* and that were modelled with the DCM. To test this hypothesis, we therefore performed several DCM inversions on the same data under quantitatively different priors, and compared the model evidence for each of these models.

Specifically we followed the following procedure:

1. We first wanted to identify which synaptic parameters we would expect to be altered in the NMDAR-Ab *in vivo* experiments, based on the microscale *in vitro* recordings. To be able to make quantitative predictions about such parameters we quantified relevant features of synaptic transmission in whole-cell patch clamp recordings. Using these features, we generated quantitative predictions of synaptic parameter differences between control-Ab and NMDAR-Ab conditions.
2. To be able to infer synaptic coupling parameters underlying the local field potential data, we extracted data from different relevant conditions and states (i.e. interictal control-Ab, interictal NMDAR-Ab, and ictal NMDAR-Ab) to then fit the microcircuit models using DCM.
3. We next wished to identify a synaptic parameter combination that best describes a physiological, interictal state from which all other conditions emerge by changes in synaptic coupling. We therefore fitted a single canonical microcircuit model to control-Ab data, using the inferred parameters as priors in subsequent Bayesian model inversions in the next steps.
4. To address the question whether the observed microscale changes in synaptic function contribute to the observed LFP features, we fitted a single canonical microcircuit to interictal NMDAR-Ab data under different sets of priors – including those that are informed by microscale data, and those that are not. This allowed us to perform Bayesian model comparison between these models. Here we would expect models that contain microscale-derived prior information to outperform other models if the modelled microscale features contribute to the LFP data.
5. To address the question which synaptic mechanisms best explain condition- and state-differences, we fitted a single canonical microcircuit to seizure NMDAR-Ab data, then quantified the parameter differences between different DCM models. The objective here was to quantify the main differences between conditions (control-Ab vs NMDAR-Ab) and states (interictal vs. seizure) in terms of neuronal population model parameters.

## Abbreviations

IgG: immunoglobulin G;
NMDAR: N-methyl D-aspartate-receptor;
Ab: antibody;
PregS: Pregnenolone sulphate

## Acknowledgments

We thank Rob Wykes, Jonathan Cornford and Andrea Lieb (University College London, UK) for use of EEG powerband analysis program. SKW was funded by an Epilepsy Research UK Fellowship (F3001) and Wellcome Trust Clinical Research Career Development Fellowship (216613/Z/19/Z) during this work. HP received support from the German Research Foundation (DFG; grant numbers PR1274/3-1, 4-1, 5-1) and the German Federal Ministry of Education and Research (BMBF; Connect-Generate). NG received support from the German Ministry for Education and Research (BMBF; 31P7398 and Connect-Generate), the National Multiple Sclerosis Society (NMSS; BRAVEinMS), the Wellcome Trust (208938/Z/17/Z) and the Forschungskommission of the Medical Faculty of the Heinrich-Heine-University.

## Contributions

SKW designed the study, carried out, analysed and interpreted data from all *in vivo* experiments, the *in vitro* whole-cell patch clamp experiments, and the *in vivo* EEG experiments. SKW performed the immunofluorescence experiments. SKW drafted the manuscript. RER helped design the study, analysed and interpreted all the dynamic causal modelling experiments, and drafted the manuscript. MAW acquired, analysed, and interpreted the *in vitro* LFP experimental data and drafted the manuscript. MAU analysed and interpreted the *in vitro* LFP experimental data and videos of experimental subjects, and also drafted the manuscript. DRD and CCB acquired, analysed and interpreted the immunofluorescence data. Additionally, DRD helped draft the manuscript. TTW analysed and interpreted the videos of experimental subjects and helped draft the manuscript. SB and NG produced, verified and provided NMDAR-Ab and control antibody preparations for experimental use. JK and HP produced, verified and provided NMDAR-Ab and control antibody preparations for experimental use. LJ produced and verified NMDAR-Ab and control immunoglobulins for experimental use. DSB supervised the dynamic causal modelling experiments and drafted the manuscript. AV designed the study, supervised SKW and drafted the manuscript. SDG supervised SKW with *in vivo* experiments, analysed and interpreted data from *in vivo* EEG and *in vitro* electrophysiology experiments. GLW designed the study, provided supervision and assistance to SKW with *in vitro* experiments, analysed and interpreted data from *in vitro* electrophysiology experiments, and drafted the manuscript. All authors substantively revised and approved the submitted version.

## Competing interests

The authors have no competing interest to declare.

## Materials and Correspondence

Requests should be addressed to Professor Gavin Woodhall.

## Supplementary Methods

### Dynamic causal modeling

#### Feature extraction from whole-cell patch clamp recordings

We quantified amplitude, decay time, and variability of sEPSCs measured from the whole-cell hippocampal patch clamp recordings described above. We selected 10 minutes of continuous, artefact free recordings from both control-Ab and NMDAR-Ab conditions, and bandpass filtered these epochs between 0.3Hz and 2000Hz. In these traces, we identified peaks with a local prominence of at least 20mV (n=1624 in control-Ab condition; n = 374 in NMDAR-Ab condition), and segmented windows of 30ms prior to the peak, and 60ms following the peak, of which we considered the first 20ms as baseline. We normalised each segment to the baseline mean and extracted the mean amplitude of these normalised peaks. For each segment we identified the time elapsed from the peak to when the signal had decayed to half-peak values, and we then calculated the mean of this period as the time constant of sEPSCs. To quantify variance in the responses, we calculated the cumulative frequencies of sEPSCs of given amplitudes (normalized to the maximal amplitude observed in each condition, respectively). We then quantified the gradient of this distribution at the 50^th^ centile to approximate the population parameter σ used in DCM. Log-differences between these average quantitative features in the control-Ab and NMDAR-Ab conditions from the whole-cell patch clamp recordings were subsequently used to inform priors of DCM inversions of LFP data.

#### LFP segmentation and data extraction

We selected six 1 hour segments for 2 animals treated with control IgG, and three 1 hour segments for 1 treated with NMDAR-Ab IgG, at the peak of the observed epileptogenesis (that is, at 48h). Each trace was z-scored and sections exceeding an absolute z-score of 5.5 were coded as seizure; else they were coded as interictal. We then added the 45s before and after the automatically identified seizure-segments to the seizure segments to capture seizure onset and seizure offset transitions. This process served to automate the LFP ictal/interictal classification confirmed by visual analysis, and we then divided all segments into 45s sections for subsequent DCM analysis. In total this process resulted in 514 control-Ab interictal segments, 0 control-Ab seizure segments, 114 NMDAR-Ab interictal segments, and 228 NMDAR-Ab seizure segments after exclusion of artefact. For each of the LFP traces, average power spectra were estimated using a multivariate autoregressive model implemented in the DCM software (40).

#### DCM fit to control data

Assuming that data recorded from control-Ab injected mice are representative of the ‘baseline’ state, we first fitted a single canonical microcircuit model (CMC)(41) using standard DCM inversion techniques (EM algorithm performing gradient descent on a free energy approximation of the negative log likelihood). This process provides us with posterior densities over neuronal parameters for the CMC (as summarized in Table 1).

#### DCM fit to NMDAR-Ab interictal data

In order to test whether the synaptic changes identified in the whole-cell patch clamp recordings contribute to the LFP features recorded in NMDAR-Ab injected mice, we fitted single CMC models under different prior parameter sets to the NMDAR-Ab interictal LFP data and compared their relative evidence. The null model used control-Ab derived parameter values exactly as priors, without any additional changes. The remainder of the model space was defined by altering prior parameter values quantitatively based on the expected control-Ab *vs*. NMDAR-Ab differences derived from the whole-cell patch clamp recordings. Specifically, we added the log difference between NMDAR-Ab and control-Ab sEPSC amplitude to excitatory coupling parameters γ1-3; the log difference between NMDAR-Ab and control-Ab sEPSC half-life to the excitatory time constants τ2,3; and the log difference in cumulative variance between NMDAR-Ab and control-Ab size distribution to the population variance parameter σ. We divided the model space to compare models where only subsets of these changes were made on the priors. These were divided along two main design features: parameter type (γ, τ, σ, or their combinations) and location in the microcircuit (superficial, fast oscillator pair; deep, slow oscillator pair; or both), resulting in a total of 7 x 3 models that carried some microscale information in their priors; and one null model without the microscale information. Comparison between these models was then made based on the free energy approximation of their respective model evidence (42). The winning model was used for subsequent analyses.

#### DCM fit to NMDAR-Ab seizure data

To identify parameter changes associated with the transition into epileptic seizures, we fitted a single CMC to NMDAR-Ab seizure LFP segments using parameters from the winning model identified in the previous step. We considered the absolute parameter differences between the DCM fit to NMDAR-Ab interictal data, and the DCM fit to NMDAR-Ab seizure data as the parameter change associated with ictogenesis for the subsequent simulations.

### Simulations

The parameters inferred by fitting the microcircuit models to LFP data are fully generative and can therefore be used to simulate novel data. We exploited this ‘simulation mode’ to test the effects of different gradual changes in parameters on the simulated LFP power spectrum. We ran simulations along two sets of parameter changes: ‘epileptogenesis’ parameters θ_E_ (i.e. the difference between control-Ab and NMDAR-Ab interictal estimated parameters); and ‘ictogenesis’ parameters θ_I_ (i.e. the difference between MMDAR-Ab interictal and NMDAR-Ab seizure estimated parameters). Simulations were in *n* = 200 steps along each of those axis, so that 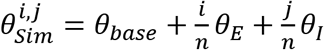 with θ_base_ representing the empirical parameters inferred from the control-Ab model inversion.

We classified each simulated power spectrum across this parameter space based on the least squares difference to any of the three empirically observed states into control-Ab-like, NMDAR-Ab interictal-like, and NMDAR-Ab seizure-like states. To quantify local changes induced by small changes in ictogenicity, we calculated the mean squared difference between (i) the power spectra derived at a given simulation parameterization, and (ii) the power spectra derived at neighbouring parameterizations along the θ_I_ direction.

**Supplementary Figure 1.**
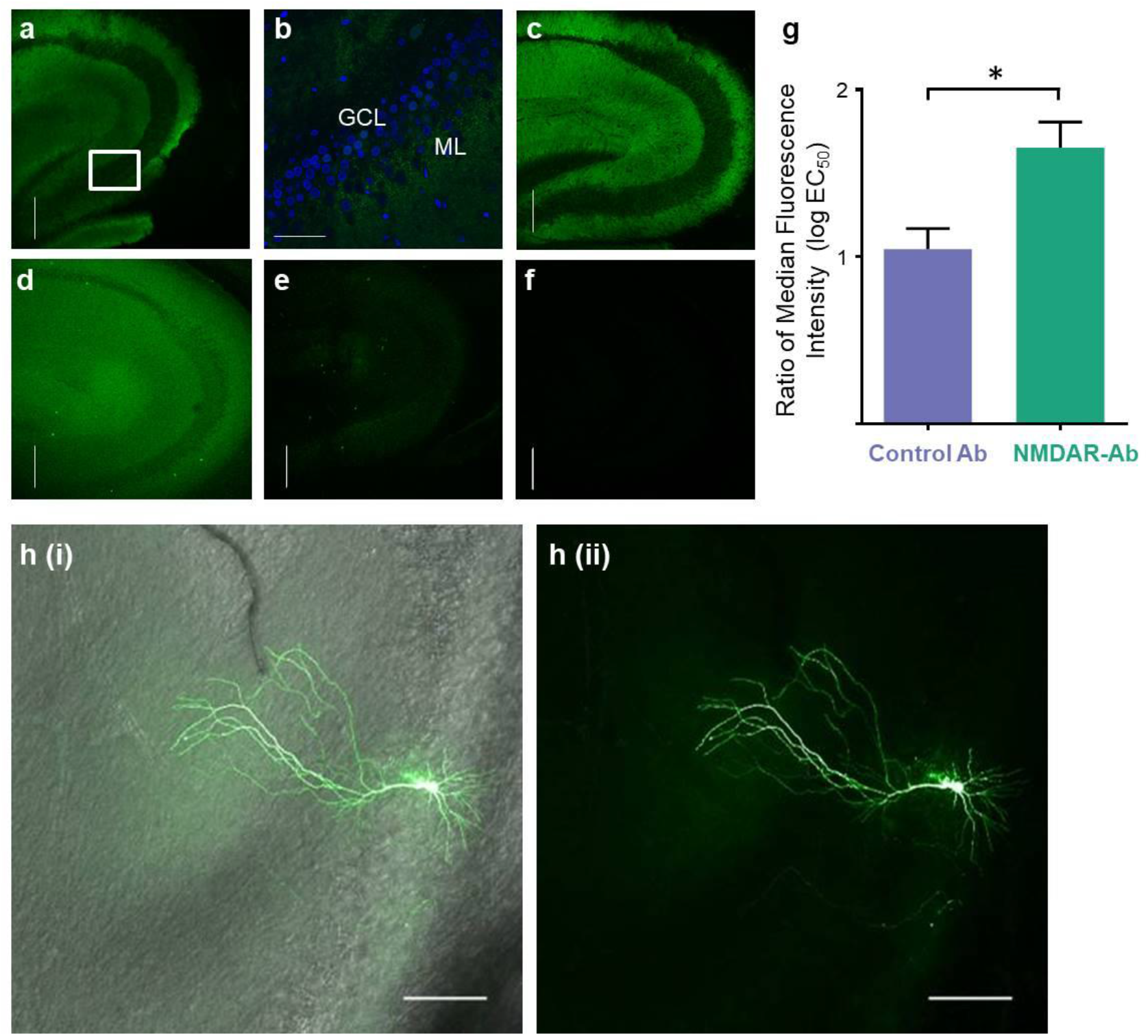
Immunohistochemistry confirming NMDAR-Ab binding to juvenile Wistar rat hippocampus and morphology of pyramidal cells in CA3 used for whole-cell patch clamp recordings. **a** Representative confocal image of hippocampus from sagittal brain slice prepared after acute ICV injection of monoclonal NMDAR-Ab SSM5 shows typical staining pattern with secondary anti-human IgG (green). Scale bar = 250μm. **b** Magnification of panel (**a**) shows the typical binding pattern of NMDAR-Abs with relative sparing of the granular cell layer (GCL) compared to the molecular cell layer (ML). Scale bar = 50μm. **c** Representative confocal image of hippocampus from sagittal brain slices prepared after chronic infusion of monoclonal NMDAR-Ab 12D7 after application of secondary anti-human IgG (green). Scale bar = 250μm. **d** Representative confocal image of hippocampus from sagittal brain slice prepared after chronic infusion of NMDAR-Ab positive IgG also shows typical staining pattern with secondary anti-human IgG (green). Scale bar = 250μm. **e** Representative confocal image of hippocampus from sagittal brain slices prepared after chronic infusion of healthy control IgG after application of secondary anti-human IgG (green). There is no specific binding of these antibodies. Scale bar = 250μm. **f** Representative confocal image of hippocampus from sagittal brain slices prepared after acute injection control monoclonal antibody mG053 after application of secondary anti-human IgG (green). There is no specific binding of these antibodies. Scale bar = 250μm. **g** The NMDAR-Ab treated brain slices showed increased median florescent intensity log EC_50_ ratios (GCL vs. ML) compared to controls (control group contains: healthy control IgG n=1 animal, 3 brain slices and control monoclonal 12D7 n=3 animals, 9 brain slices; NMDAR group contains: SSM5 monoclonal n=8 animals, 22 brain slices, 003-102 n=2 animals, 2 brain slices and patient IgG n=3 animals 5 brain slices; p=0.04, Mann-Whitney). **h** Cells used for spontaneous excitatory and inhibitory current recordings were from putative pyramidal cells with the CA3 region; an example cell injected with neurobiotin shows clear features of a pyramidal cell (green, anti-streptavidin IgG). Scale bar 150 μm.

**Supplementary Figure 2.**
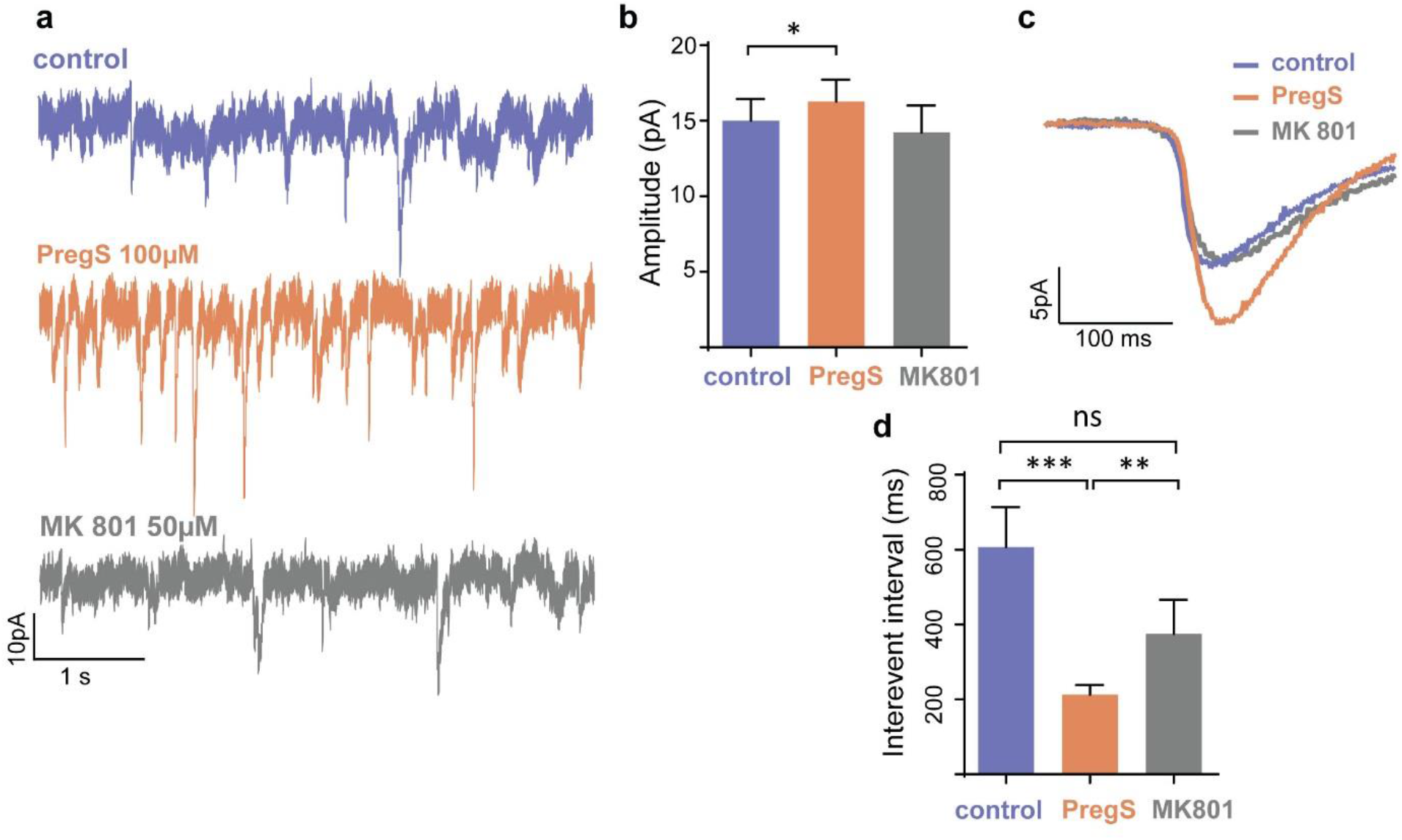
Pregnenolone sulphate increases synaptic levels of NMDARs in naïve rat hippocampal neurons and glutamatergic neurotransmission in rat brain slices *in vitro*. **a** Representative whole-cell patch clamp sEPSC recordings from CA3 pyramidal cells in untreated (no antibody) hippocampal slices. Scale bar 10pA vs. 1s. **b** Amplitude (pA) of sEPCS recordings from putative CA3 pyramidal neurons before and after addition of PregS (n=16 cells; p= 0.04, Wilcoxon-paired rank test); and of MK801 (n=9 cells; p=0.1, Wilcoxon-paired rank test). **c** Representative averaged amplitude (pA) of sEPSCs after addition of PregS and MK801 Scale bar 5pA vs. 100ms. **d** Interevent interval (ms) of sEPSCS recordings from putative CA3 pyramidal neurons before (control) and after addition of PregS (n=20; p=<0.001, Wilcoxon paired rank test) and of MK801 (n=9; p=0.01, Wilcoxon paired rank test).

**Supplementary Table 1.** Details of antibodies used in each experiment.

| Procedure | Number of animals used with specified antibody |  |  |  |  |  |
| --- | --- | --- | --- | --- | --- | --- |
|  | Healthy control IgG | Control monoclonal antibody mG053 | Control monoclonal antibody 12D7 | NMDAR monoclonal antibody SSM5 | NMDAR monoclonal antibody 003-102 | NMDAR -Ab patient IgG |
| ICV injection | N=3/6 | - | N=3/6 | N=8/11 | - | N=3/11 |
| Osmotic pump infusion with EEG transmitter | N=3/6 | N=3/6 | - | N=2/6 | N=2/6 | N=2/6 |

ICV intraventricular injection NMDAR N-methyl-D-aspartate receptor Ab antibody

**Supplementary Table 2.** Patient details for human tissue electrophysiology experiments.

| Patient | Operation | Medication | Histology |
| --- | --- | --- | --- |
| 1 | Frontal resection | Topiramate<br>Clobazam | Focal cortical dysplasia Type IIB |
| 2 | Frontal resection | Lamotrigine<br>Carbamazepine<br>Clobazam | Focal cortical dysplasia Type IB |
| 3 | Frontal resection | Modified KD<br>Sodium valproate<br>Vigabatrin | No definite abnormal histology |
| 4 | Temporal parietal occipital disconnection | Topiramate<br>Clobazam | No definite evidence of epileptogenic lesion |
| 5 | Temporal lobectomy | Lamotrigine | Changes consistent with epilepsy |
| 6 | Temporal lobectomy | Carbamazepine<br>Levetiracetam | Hippocampal sclerosis |
| 7 | Right hemispherotomy | Carbamazepine<br>Levetiracetam<br>Clonazepam | Hippocampal sclerosis |

### Supplementary videos

**Video 1** Hyperexcitable phenotype captured 48 hours after NMDAR-Ab ICV injection in juvenile Wistar rats.

**Video 2a-c** Example video clips of epileptic activity in NMDAR-Ab chronically infused juvenile Wistar rats during wakefulness (a, b), and in sleep (c).

## Notes

### Competing Interest Statement

The authors have declared no competing interest.

